# Metagenomic insight into the plant growth-promoting potential of a diesel-degrading bacterial consortium for enhanced rhizoremediation application

**DOI:** 10.1101/2021.03.26.437261

**Authors:** Michael O. Eze, Volker Thiel, Grant C. Hose, Simon C. George, Rolf Daniel

**Author notes:** These authors contributed equally to this work (Simon C. George, Rolf Daniel).

## Abstract

The slow rate of natural attenuation of organic pollutants, together with unwanted environmental impacts of traditional remediation strategies, has necessitated the exploration of plant-microbe systems for enhanced bioremediation applications. The identification of microorganisms capable of promoting both plant growth and hydrocarbon degradation is crucial to the success of plant-based remediation techniques. Through successive enrichments of a soil sample from a historic oil-contaminated site in Wietze, Germany, we isolated a plant growth-promoting and hydrocarbon-degrading bacterial consortium. Metagenome analysis of the consortium led to the identification of genes and taxa putatively associated with these processes. The majority of the coding DNA sequences involved in these reactions were affiliated to *Acidocella aminolytica* and *Acidobacterium capsulatum.* In microcosm experiments performed in association with *Medicago sativa* L., the consortium achieved 91% rhizodegradation of diesel fuel hydrocarbons within 60 days, indicating its potential for biotechnological applications in the remediation of sites contaminated by organic pollutants.

## INTRODUCTION

Phytoremediation is the remediation of contaminants by plant-based techniques. This approach offers an environmentally friendly and cost-effective in-situ method for the remediation of contaminated soils ^1^, and has been applied to both organic and inorganic contaminants. Closely related to phytoremediation is rhizoremediation, which is the degradation of contaminants by root-associated microorganisms. To enhance the effectiveness of phytoremediation and rhizoremediation, plant growth-promoting rhizobacteria (PGPR) have been the focus of research in recent decades ^2,3^

PGPR inhabit the rhizospheric zones of plants and can directly or indirectly influence plant growth. PGPR provide nutrients to host plants, produce phytohormones that regulate plant growth and metabolic activities, and protect host plants from pathogens and abiotic stress ^4,5^. The plant growth-promoting activities of PGPR include nitrogen fixation, phosphate and potassium solubilization, indoleacetic acid and pyrroloquinoline quinone biosynthesis, siderophore transport, induction of systemic resistance, and interference with pathogen toxin production ^6–8^.

PGPR readily establish in soils due to their high growth rate and adaptability to a wide variety of environments, and ability to metabolize a wide range of natural and xenobiotic compounds ^6^. Consequently, there is an increasing interest in enhancing rhizoremediation through the inoculation of microbial consortia with the required metabolic pathways ^9,10^. Unfortunately, research to date has focused just on hydrocarbon-degrading microbes ^11,12^, with very few studies targeting organisms capable of both plant growth promotion and hydrocarbon degradation ^13^.

Rhizospheric soils derived from petroleum contaminated sites are often a rich source of microorganisms that have the metabolic ability to degrade organic contaminants while enhancing plant growth ^14–16^. Since microorganisms present in oil-polluted sites often possess adaptability and resistance to toxic organic compounds, an examination of their plant growth-promoting ability will help bridge the knowledge gap required to develop effective plant growth-promoting inocula for plants in contaminated soils. The application of such inocula will enhance the tolerance of plants to hydrocarbon toxicity, promote biomass production, and enhance rhizoremediation. Thus, the aims of this study were to isolate a bacterial consortium capable of enhancing plant growth and hydrocarbon degradation. In microcosm experiments performed in association with *Medicago sativa* L., the consortium significantly enhanced the growth of *M. sativa*, and achieved 91% rhizodegradation of diesel fuel hydrocarbons within 60 days. The choice of *M. sativa* was based on our previous study that revealed the plant’s tolerance to hydrocarbons ^17^. The results of this study will expand the range of available PGPR for use in rhizoremediation of environmental contaminants.

## RESULTS

### Bacterial diversity in the consortium

The consortium used in this study was isolated by Eze, et al. ^18^ from a crude-oil polluted site in Wietze, Germany. The bacterial diversity in both the original soil sample and the enrichment culture was discussed in Eze, et al. ^18^.

The relative abundance of classifiable bacterial sequences based on metagenome analysis showed the dominance of Alphaproteobacteria in the original soil sample, with relative abundance of 22% (Figures 1). This was followed by Acidobacteriia (9%), Betaproteobacteria (8%), Gammaproteobacteria (6%), Actinobacteria (5%), Deltaproteobacteria (2%), Sphingobacteria (1%) and Planctomycetia (1%). Viruses and Archaea accounted for 0.09% and 0.3% of the total sequences, while Eukaryota accounted for 3% (of which 2% are Fungi). At genus level, *Bradyrhizobium* (3%) dominated in the original soil followed by *Pseudomonas* (2%). Other genera present included *Sphingomonas, Mycobacterium, Mucilaginibacter, Acidocella, Acidobacterium* and *Aquabacter.* The successive enrichments resulted in a consortium with approximately 60% relative abundance of Alphaproteobacteria (predominantly *Acidocella),* followed by Acidobacteriia (Figures 1b and 1c). Other bacterial classes with representative abundance (at least 1%) in the consortium include Betaproteobacteria, Gammaproteobacteria and Actinobacteria. At the genus level, *Acidocella, Acidobacterium* and *Acidiphilium* dominated (Figure 1b).

**Figure 1.**
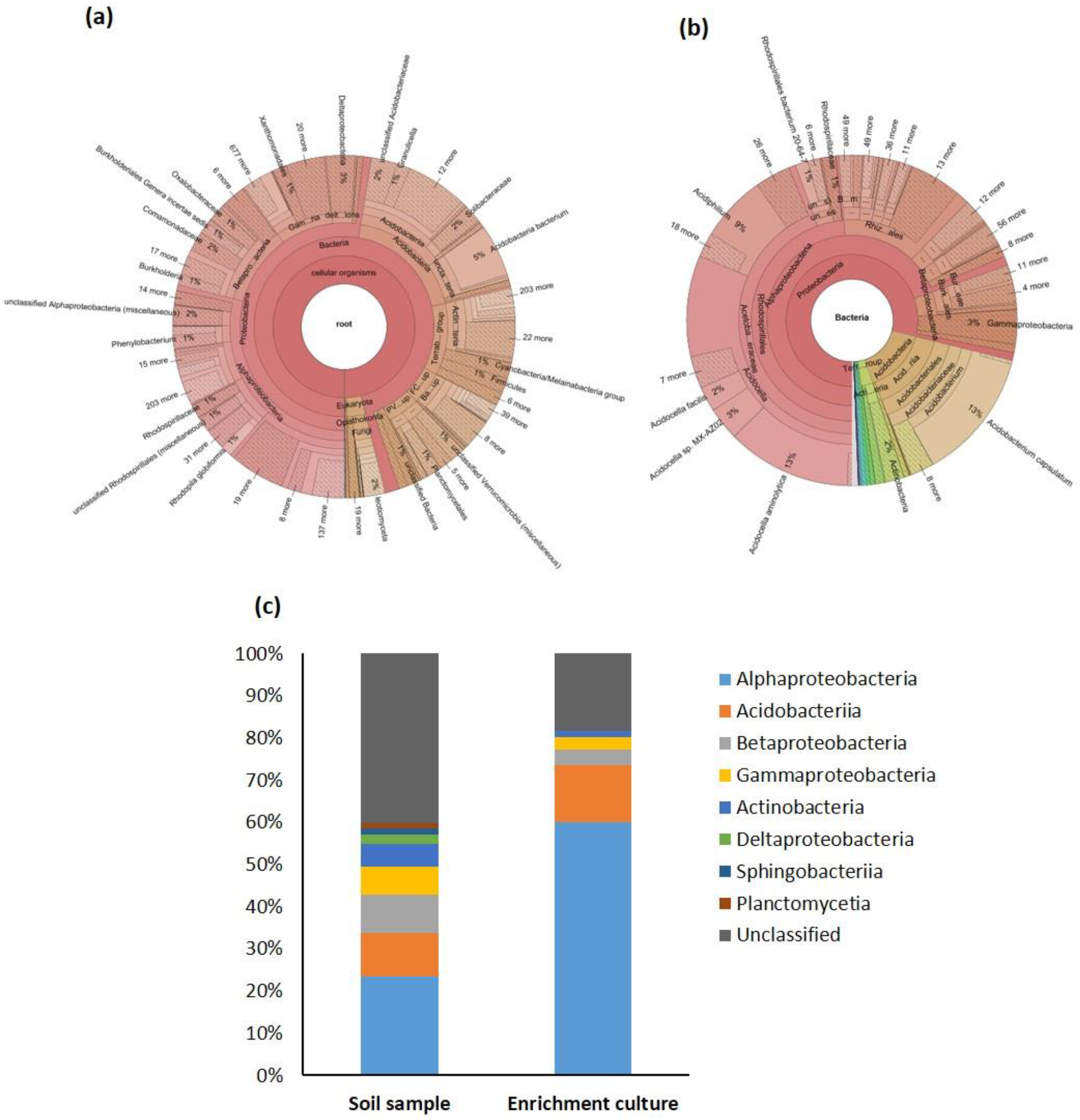
Taxonomic classification of (a) the original soil sample and (b) the enrichment culture based on metagenome data, with (c) a summary column chart of the key differences in the bacterial community composition.

### Identification of plant growth-promoting enzymes

Functional analysis of the bacterial consortium revealed the presence of 26 enzymatic classes involved in plant growth-promoting activities putatively (Figure 2). The majority of the 177 coding DNA sequences (CDSs) potentially involved in these reactions were associated with phosphate solubilization and nitrogen fixation (99 and 55 CDSs, respectively). These were followed by pyrroloquinoline quinone synthesis (13 CDSs), zinc solubilization (5 CDSs), siderophore transport (3 CDSs), and indoleacetic acid synthesis (2 CDSs).

**Figure 2.**
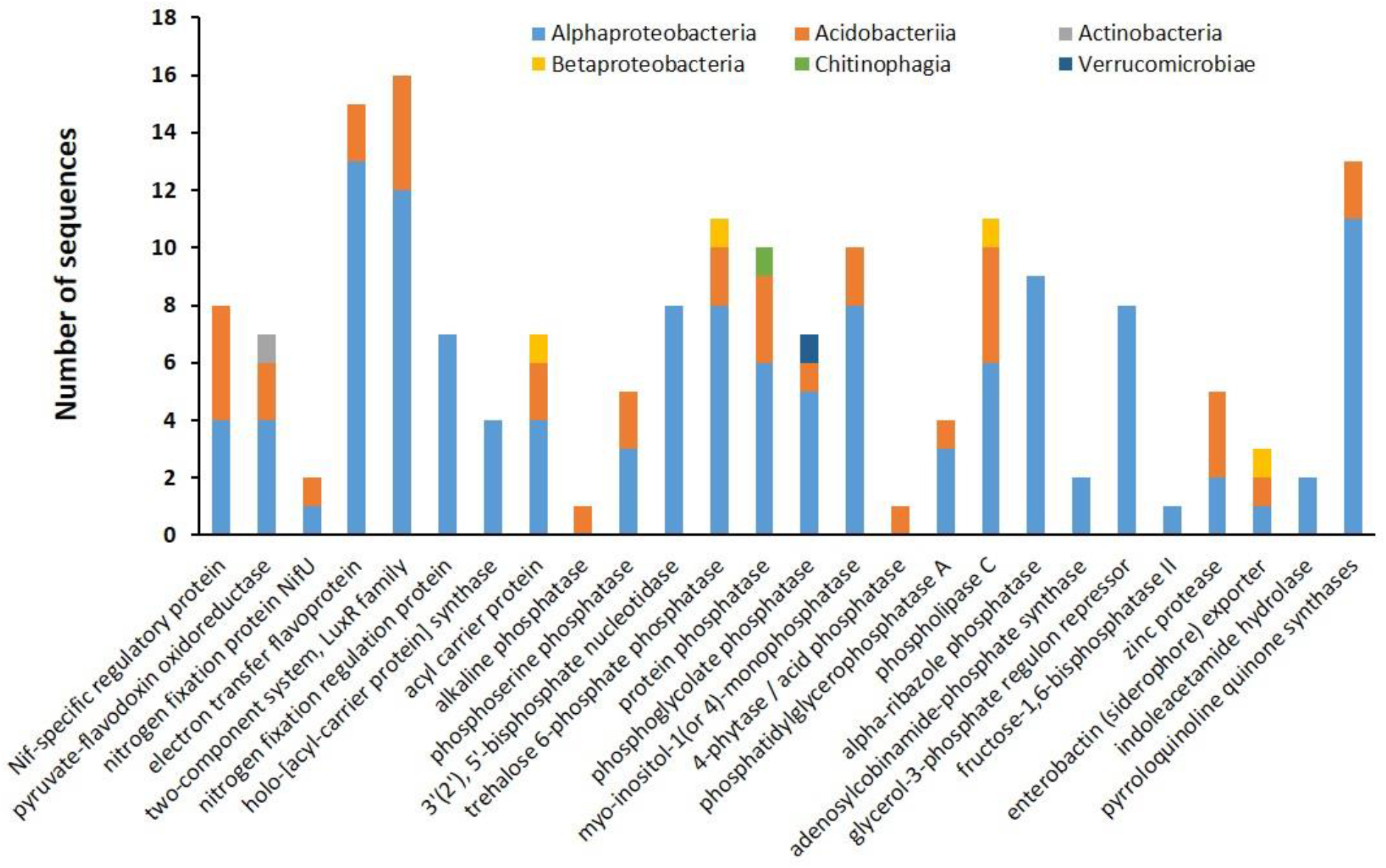
Key enzymes involved in plant growth-promoting activities and their taxonomic assignation at the class level. For detailed information, see Supplementary Table S1.

### Taxonomic assignation of CDSs associated with nitrogen uptake

Functional analysis of the metagenome data revealed the presence of 55 CDSs potentially involved in nitrogen uptake by plants. These include the *nifAJU* and *fixABJKL* genes ^19–21^. Taxonomic assignment revealed that 56% of the CDSs involved in nitrogen uptake (31 CDSs) belonged to the *Acidocella* and *Acidobacterium* genera (Figure 3). Other represented genera include *Acidiphilium, Rhodopseudomonas, Acetobacter, Asaia* and *Bradyrhizobium.* At the species level, *Acidocella aminolytica* accounted for the majority of the CDSs assigned to *Acidocella* (Supplementary Table S1). Other represented genera included *Acidiphilium* (3 CDSs); *Acetobacter, Asaia* and *Rhodopseudomonas* (with 2 CDSs each). Six CDSs could not be assigned to any specific bacterial genus.

**Figure 3.**
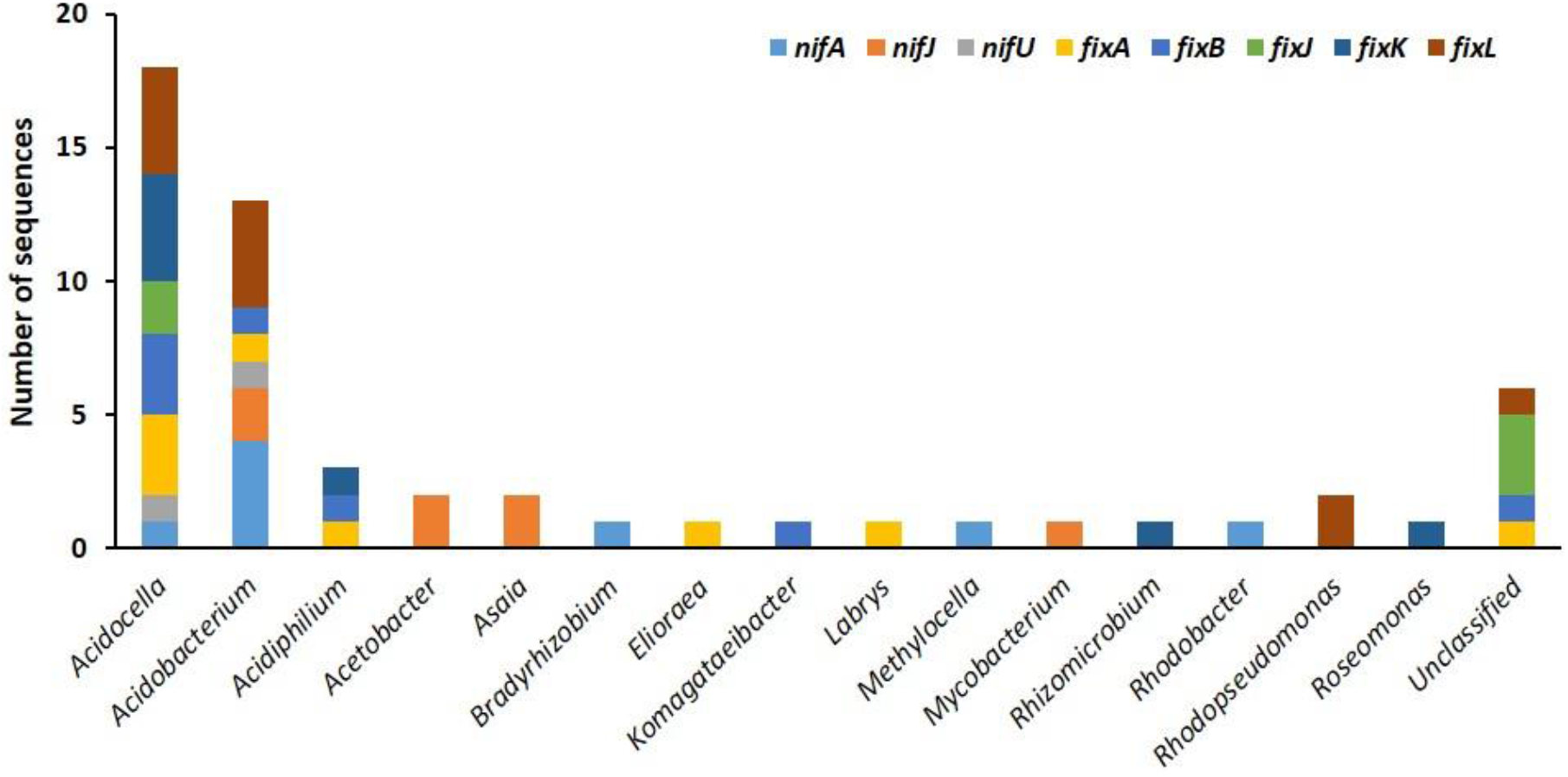
Genus assignment of the genes involved in biological nitrogen fixation. *nifA:* Nif-specific regulatory protein; *nifJ:* pyruvate-flavodoxin oxidoreductase; *nifU:* nitrogen fixation protein NifU; *fixA:* electron transfer flavoprotein beta subunit; *fixB:* electron transfer flavoprotein alpha subunit; *fixJ:* two-component system, LuxR family, response regulator; *fixK:* nitrogen fixation regulation protein; *fixL:* two-component system, LuxR family, sensor kinase.

### Taxonomic assignation of CDSs associated with phosphate solubilization

Bacterial taxa involved in phosphate solubilization include Alphaproteobacteria such as *Acidocella, Acidiphilium, Methylovirgula, Roseovarius, Skermanella* and *Sphingomonas;* Betaproteobacteria such as *Paraburkholderia* and *Hydrogenophaga;* Acidobacteriia such as *Acidobacterium capsulatum;* Chitinophagia such as *Niastella;* and Verrucomicrobiae such as *Pedosphaera* (Supplementary Table S1). The majority of the CDSs belonged to the *Acidocella* (68%) and *Acidobacterium* (19%). At the species level, the majority of the CDSs belonged to *Acidocella aminolytica* (44 CDSs) and *Acidobacterium capsulatum* (19 CDSs) (Supplementary Table S1).

### Zinc solubilization, siderophore, indoleacetic acid and pyrroloquinoline quinone syntheses

The results of the metagenome analysis showed that 5 CDSs were putatively involved in zinc solubilization, 3 CDSs in siderophore transport, 2 CDSs in indoleacetic acid synthesis, and 13 CDSs in pyrroloquinoline quinone synthesis (Figure 4). Of the five zinc-solubilizing CDSs, three were assigned to *Acidobacterium capsulatum,* one to *Acidocella aminolytica* and one to *Acidocella facilis* (Supplementary Table S1). The three CDSs involved in siderophore production were assigned to *Acidobacterium, Acidocella* and an unclassified *Burkholderiales* (1 CDS each) (Figure 4a). The two CDSs responsible for indoleacetic acid synthesis were assigned to *Hyphomicrobium* and *Mesorhizobium* genera (Figure 4b). The majority of the pyrroloquinoline quinone synthases belong to *Acidocella* (9 CDSs), followed by *Acidobacterium* (2 CDSs), *Acidiphilium* (1 CDS) and *Bosea* (1 CDS).

**Figure 4.**
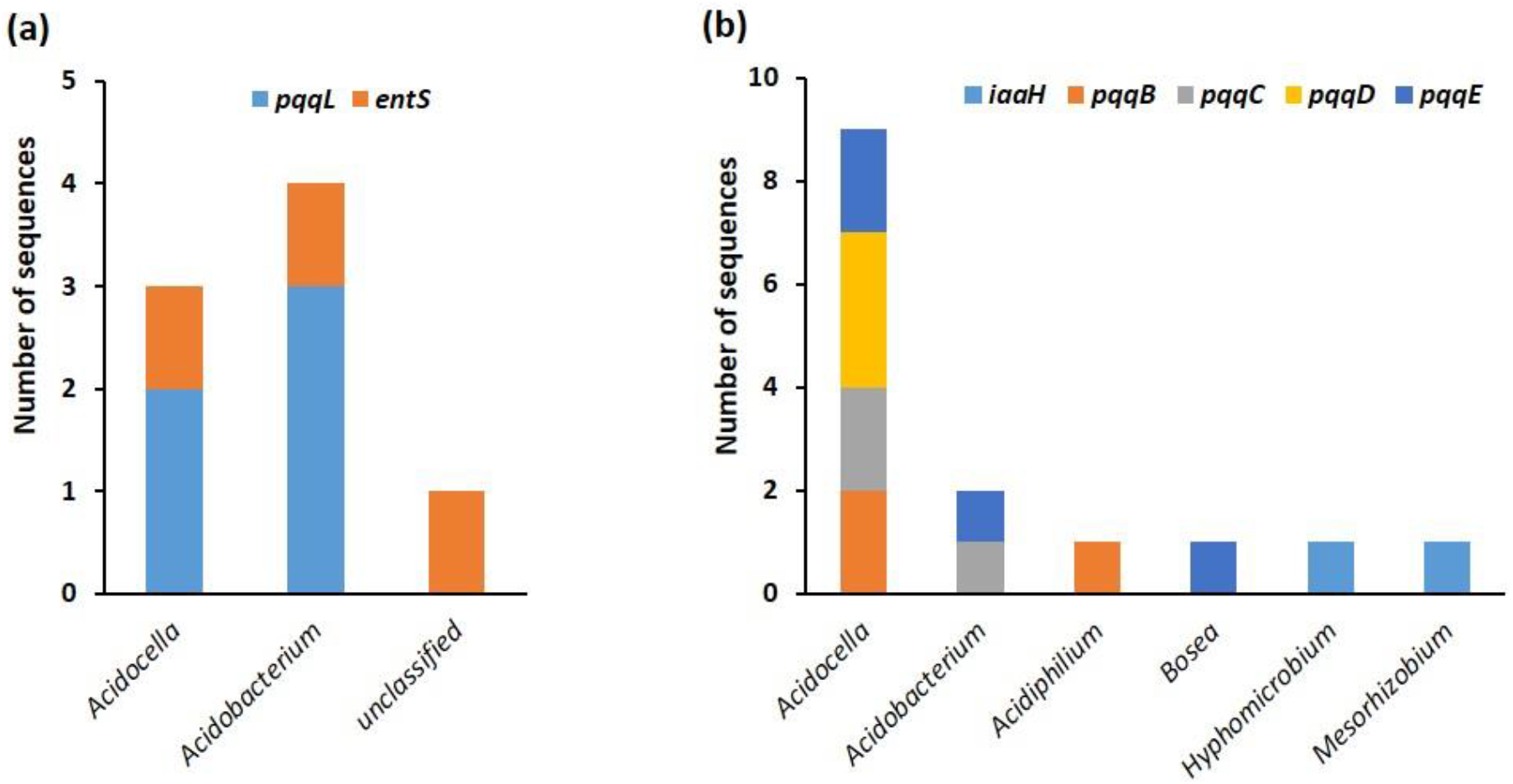
Genus assignment of the genes involved in (a) zinc solubilization and siderophore transport; and (b) indoleacetic acid and pyrroloquinoline quinone biosynthesis. *pqqL*: zinc protease; *entS:* Enterobactin (siderophore) exporter; *iaaH*: indoleacetamide hydrolase; *pqqB*: pyrroloquinoline quinone biosynthesis protein B; *pqqC*: pyrroloquinoline-quinone synthase; *pqqD*: pyrroloquinoline quinone biosynthesis protein D; *pqqE*: PqqA peptide cyclase.

### Effect of bacterial inoculation on plant growth and biomass production

The inoculation of *Medicago sativa* with the isolated consortium resulted in increased growth and biomass production of the inoculated plants (Figure 5). The mean dry biomass (± standard error) of *M. sativa* plants inoculated with the consortium was 5.48 ± 0.07 g, significantly greater (*p* < 0.001) than the dry biomass of plants in the uninoculated control (3.30 ± 0.07 g).

**Figure 5.**
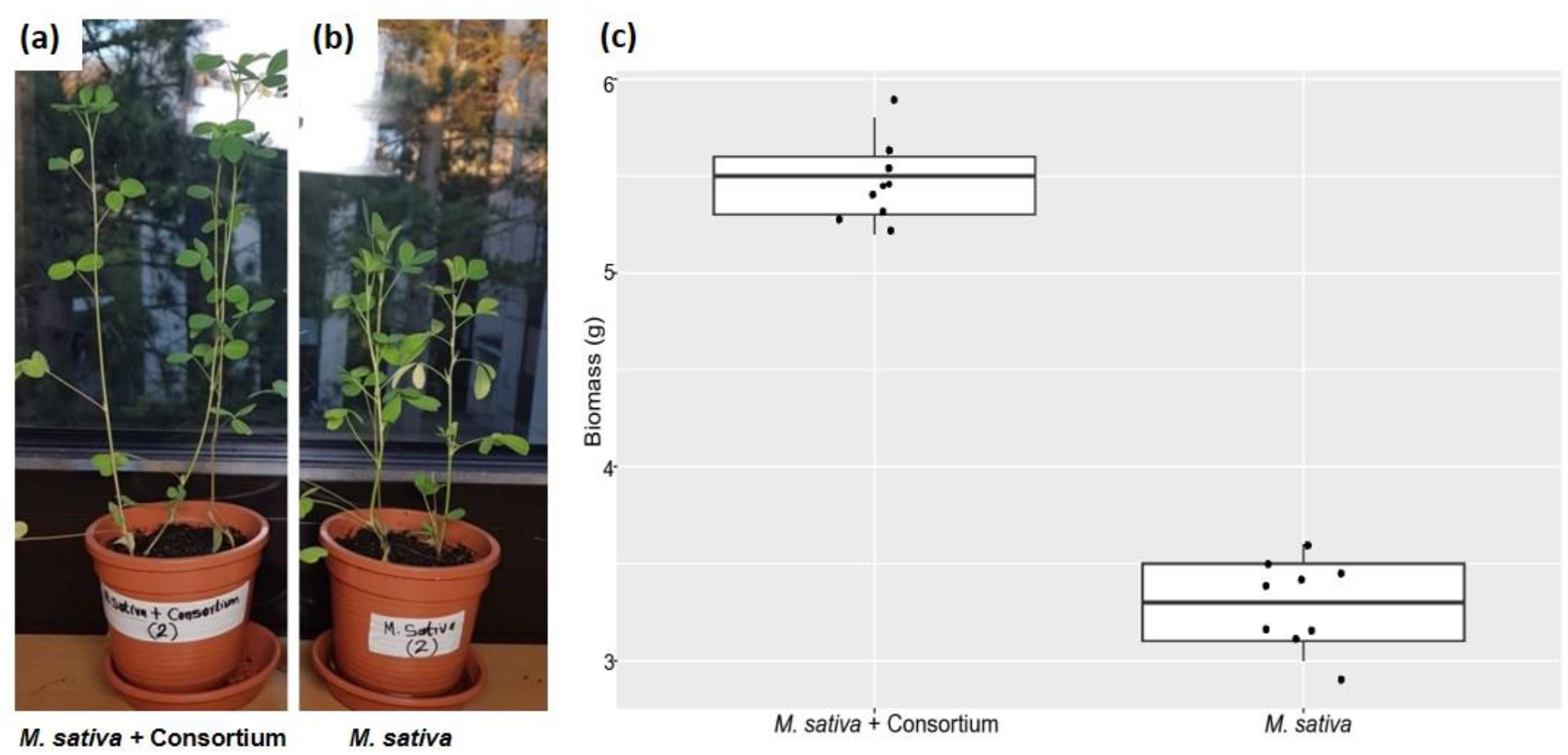
Changes in growth and biomass production of (a) *Medicago sativa* plants inoculated with the isolated microbial consortium, in comparison with (b) uninoculated *Medicago sativa* plants. (c) Boxplot showing significant differences in biomass production of inoculated and uninoculated plants.

### Effect of inoculation on biodegradation of petroleum hydrocarbons

Microbial inoculation of *M. sativa* enhanced the biodegradation of petroleum hydrocarbons. The mean residual total petroleum hydrocarbon concentrations (± standard error) for the “Control” (representing natural attenuation) was 2.77 ± 0.03 g/Kg, for “*M. sativa*” was 1.36 ± 0.04 g/Kg, for the “Consortium” was 1.15 ± 0.05 g/Kg, and for *“M. sativa* + Consortium” was 0.40 ± 0.03 g/Kg (Figure 6a). These concentrations represent 40%, 70%, 75% and 91% degradation, respectively. Statistical analysis showed that the effects of the different treatment options were significantly different *(p* < 0.05) from each other. Gas chromatography-mass spectrometry (GC-MS) of solvent extracts prepared from the experimental soils revealed a near complete degradation of both the aliphatic and aromatic hydrocarbons in diesel fuel in the *“M. sativa* + Consortium” treatment (Figure 6b and 6c). Thus, while natural attenuation resulted in the lowest degradation efficiency, synergistic interactions of *M. sativa* and the consortium led to a significantly higher degradation efficiency.

**Figure 6.**
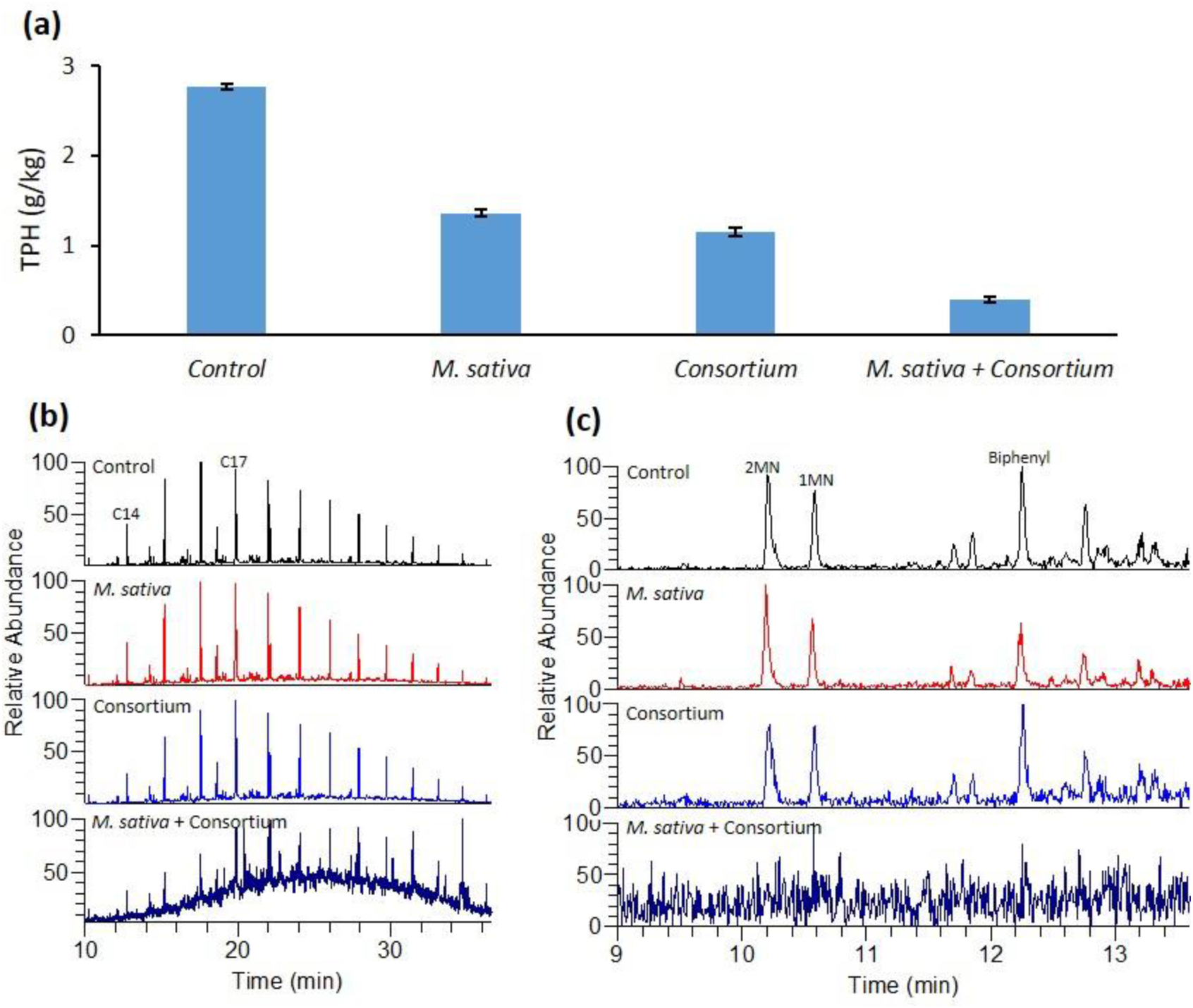
(a) Bar chart showing the mean values of residual total petroleum hydrocarbons (TPH, g/Kg) for the different treatments after 60 days (n=3). (b) Partial *m/z* 57 mass chromatograms showing a homologous series of residual *n*-alkanes in the contaminated soils under different treatments (carbon numbers annotated for *n*-tetradecane and *n*-heptadecane). (c) Partial *m/z* 142 + 154 mass chromatograms of residual soils at the end of the experiment showing differential degradation of methylnaphthalene isomers (MN) and biphenyl under different treatments.

## DISCUSSION

The use of plants to clean up contaminated sites is an eco-friendly and cost-effective technology. However, petroleum hydrocarbons are phytotoxic to most plant species, and consequently impact negatively on plant growth and biomass production ^17^. Most plants are unable to prosper in contaminated soils due to hydrocarbon-induced toxicity. Therefore, the isolation of microbial consortia capable of stimulating plant growth and enhancing contaminant degradation in the rhizosphere is crucial to the success of plant-based remediation techniques. The bacterial consortium cultured in this study enabled both enhanced growth of *M. sativa* and the degradation of hydrocarbons added to the soil.

The successive enrichment of microorganisms in the contaminated soil resulted in a shift in microbial community composition and diversity. Although *Alphaproteobacteria* were dominant in both the original soil sample and the enrichment culture, successive enrichment using diesel fuel resulted in an increase in the percentage of Alphaproteobacteria from 22% in the soil sample to approximately 60% in the enrichment culture, with *Acidocella* being the dominant genus. Similarly, previous studies have shown that with increasing duration of hydrocarbon contamination in the environment, there is usually a preferential increase in Alphaproteobacteria over Gammaproteobacteria ^22^.

The consortium isolated from this study contains enzymes involved in nitrogen uptake by plants. These include the *nif* and *fix* genes. Studies of nitrogenases have revealed that they require electrons to reduce N_2_, and either a flavodoxin or a ferredoxin is required to transfer electrons from a donor to a nitrogenase, and that *nifF* and *nifJ* are responsible for such electron transport ^19,23^. In addition, *nifA* encode a number of regulatory proteins involved in nitrogen fixation ^24^. Besides the *nif* genes, regulation of nitrogen fixation is dependent on the *fix* genes ^21^. The majority of CDSs putatively encoding for nitrogen uptake in the metagenome data were assigned to *Acidobacterium* and *Acidocella*. This is not surprising considering the dominance of these genera in the consortium (Figure 1). A recent study of various strains of *Acidobacteria* revealed that they harboured homologous genes for nitrogen uptake ^25^. The presence of similar genes in the consortium indicate its potential to enhance nitrogen uptake. Nitrogen is an important limiting element for plant growth and production. It is required for the synthesis of macromolecules such as amino acids, nucleic acids, and chlorophyll. Significantly, efforts toward maximizing plant’s nitrogen uptake, translocation and assimilation by rhizobacteria inoculation have attracted much attention in recent decades ^26–30^.

Metagenome analysis revealed the presence of genes putatively responsible for phosphate solubilization in our microbial consortium (99 CDSs, Supplementary Table S1). These include *acpPS* (acyl carrier proteins), *phoAB* (alkaline phosphatase), *otsB* (trehalose 6-phosphate phosphatase), *gph* (phosphoglycolate phosphatase), and *plc* (phospholipase). Previous studies have shown that acyl carrier proteins play a crucial role in the synthesis of fatty acids, which are necessary for cellular membranes’ integrity, fluidity and permeability ^31,32^. Fatty acids also confer structural barriers to the environmental stresses that plants face ^33^. Five of the seven CDSs putatively encoding phosphoglycolate phosphatase were assigned to *Acidocella* (Supplementary Table S1). This enzyme plays a major role in photorespiratory 2-phosphoglycolate metabolism, an essential pathway for photosynthesis in plants ^34–36^. Similarly, seven of the eleven CDSs encoding trehalose 6-phosphate phosphatase, an enzyme that catalyses the biosynthesis of trehalose, were assigned to *Acidocella.* Trehalose has been found to enhance the tolerance of plants to stress ^37^. Consequently, the abundance of such phosphatases in our consortium may account for not only the growth promotion of *M. sativa* plants but also their tolerance to diesel fuel toxicity.

Functional analysis of the metagenome of the consortium from the enrichment revealed the presence of other potential genes important for plant growth promotion. These genes include *pqqL* (for zinc solubilisation), *entS* (for siderophore transport), *iaaH* (for indoleacetic acid synthesis) and *pqqBCDE* (for pyrroloquinoline quinone synthesis) (Supplementary Table S1). The enzymes that they encode are vital for plant growth and tolerance to contaminant toxicity. For example, the defence strategies of plants against pathogens has been linked to their ability to solubilize zinc ^38^. Rhizobacteria that are able to release iron-chelating molecules serve to attract iron towards the rhizosphere, where it can be absorbed by the plants ^39^. In addition, siderophore-producing PGPR can hinder the growth of pathogens by limiting the iron available for the pathogen, mostly fungi, which are unable to absorb the iron–siderophore complex ^40,41^. Previous studies on siderophore and pyrroloquinoline quinone biosynthesis have focused on *Pseudomonas* ^42^. Similarly, studies on indoleacetic acid production by PGPR have generally been limited to a few microorganisms such as *Azospirillum*, *Pseudomonas*, *Rhizobium* and *Burkholderia* ^7,43–46^. Hence, the identification of *Acidocella* and *Acidobacterium* as being potentially involved in these processes, coupled with their hydrocarbon-degrading potentials ^18^, significantly expands the range of PGPR applicable for microbially-enhanced phytoremediation of organic pollutants.

During the pot experiment, the consortium significantly enhanced plant growth and hydrocarbon degradation. The inoculation of *M. sativa* with the consortium significantly enhanced plant growth and biomass production (66%, Figure 5), indicating that the consortium was effective in promoting the growth of *M. sativa* in contaminated soils. Similarly, geochemical analyses revealed not only a reduction in the total petroleum hydrocarbons but also the degradation of both aliphatic and aromatic hydrocarbons (Figure 6). In terms of unitary application, *M. sativa* and the consortium resulted in 70% and 75% degradation respectively, indicating that the consortium was marginally more effective for hydrocarbon degradation than *M. sativa* alone. These results differed from that of a previous study by Garrido-Sanz, et al. ^10^, in which the isolated consortium was less effective than *M. sativa* for hydrocarbon remediation. The difference in remediation effectiveness between the two consortia may be attributed to differences in their bacterial community composition. The consortium isolated by Garrido-Sanz, et al. ^10^ was dominated by Gammaproteobacteria, especially *Pseudomonas.* In contrast, Alphaproteobacteria dominated in the consortium isolated in this study (Figure 1). The hydrocarbon-degrading ability of the consortium is evidently associated with the presence of CDSs putatively encoding for monooxygenase and dioxygenases. Previous metagenome analysis of the consortium revealed the presence of key monooxygenases and dioxygenases including, but not limited to, long-chain alkane monooxygenase *(ladA),* toluene monooxygenase *(tmoCF),* benzene/toluene/chlorobenzene dioxygenase *(todABC1C2)* and ethylbenzene dioxygenase *(etbAaAbAc)* ^18^. Central metabolism of aromatic hydrocarbons that follows initial activation involves ortho- and meta-cleavage of catechol or methylcatechol, and these reactions are orchestrated by enzymes such as catechol 1,2-dioxygenase and catechol 2,3-dioxygenase ^47–53^. These genes are also present in the metagenome data of the consortium ^18^.

Our experiments using the isolated consortium highlighted the role of plant-microbe synergy in the remediation of environmental pollutants. Although bioremediation by either *M. sativa* or the consortium significantly enhanced biodegradation of petroleum hydrocarbons, the greatest effectiveness was achieved in the *M. sativa* + Consortium treatment (91% degradation within 60 days). These results are of interest not only for biotechnological applications aimed at phytoremediation of toxic compounds, but also for improving crop yield in agriculture ^54,55^. The affiliation of most of the plant growth-promoting and hydrocarbon-degrading activities to the two dominant species, namely, *Acidocella aminolytica* and *Acidobacterium capsulatum,* is an indication that these species are promising candidates for biotechnological applications in the remediation of organic contaminants.

## MATERIALS AND METHODS

### Study site description

Wietze is an important historical site in terms of crude-oil production. Industrial extraction of petroleum in Germany began in 1859 ^56^. Between 1900 and 1920, Wietze was the most productive oil field in Germany, and almost 80% of German crude oil was produced here. Although oil production ceased at Wietze several decades ago, there are still a significant number of oil seeps, contaminating soils, and water bodies (Supplementary Figure S1). These sites harbour several plant species. Therefore, Wietze is an ideal place to mine for plant growth-promoting and hydrocarbon-degrading microorganisms.

### Soil sampling, enrichment and growth conditions

Approximately 10 g of topsoil (0-10 cm) was taken from a heavily polluted site located at the historical oil field at Wietze (52°39’0’’N, 09°50’0’’E), Germany in November 2019 ^18^. The sample was transported to the laboratory on ice.

A 1 g subsample of the field sample was added to an Erlenmeyer flask (300 mL) containing 100 mL of liquid mineral medium. Mineral medium was KH_2_PO_4_ (0.5 g/L), NaCl (0.5 g/L), NH_4_Cl (0.5 g/L). Sterile-filtered trace elements (1 mL/L) ^57^, vitamin solution (1 mL/L) ^57^ and MgSO_4_·7H_2_O (5 mL of a 100 mg/mL) were added post-autoclaving. Sterile-filtered diesel fuel (1 mL) was added to the flask as the sole carbon and energy source. The culture was grown at 30°C with constant shaking at 110 rpm (INFORS HT shaker, model CH-4103, Infors AG, Bottmingen, Switzerland) and subcultured every five days. After three successive enrichments, microbial cells were harvested for both metagenome studies and a greenhouse experiment.

### DNA extraction

Microbial cells (OD_600_ = 0.635) from 30 mL of the enrichment culture were harvested by centrifugation at 4000 × g for 10 min. The supernatant was subsequently discarded. DNA from the cell pellets was extracted using the PowerSoil^®^ DNA Extraction kit (Qiagen, Hilden, Germany).

### DNA extraction, metagenome sequencing, assembly and analysis

The extraction of the DNA was described in Eze, et al. ^18^. In brief, microbial cells (OD_600_ = 0.635) from 30 mL of the enrichment culture were harvested by centrifugation at 4000 × g for 10 min. DNA from the cell pellets was extracted using the PowerSoil^®^ DNA Extraction kit (Qiagen, Hilden, Germany). Sequencing libraries were generated from environmental DNA. These were barcoded using the Nextera XT-Index kit (Illumina, San Diego, USA) and the Kapa HIFI Hot Start polymerase (Kapa Biosystems, Wilmington, USA). The Göttingen Genomics Laboratory determined the sequences employing an Illumina HiSeq 2500 system using the HiSeq Rapid SBS kit V2 (2×250 bp). Metagenomic reads were further processed as described in Eze, et al. ^16^. In brief, reads were processed with the Trimmomatic tool version 0.39 ^58^ and assembled using metaSPAdes version 3.13.2 ^59^. Coverage information for each scaffold was determined using Bowtie2 version 2.3.2 ^60^ and SAMtools version 1.7 ^61^. Functional annotation of coding DNA sequences putatively encoding for the various plant growth-promoting enzymes was as described previously ^18^, and taxonomic assignment was performed using kaiju version 1.7.3 ^62^

### Potential plant growth-promoting enzymes

The enzymes of interest include nif-specific regulatory protein, pyruvate-flavodoxin oxidoreductase, nitrogen fixation protein, electron transfer flavoprotein, nitrogen fixation regulation protein, phosphatases, acyl carrier protein, phospholipases, zinc protease, enterobactin (siderophore) exporter, indoleacetamide hydrolase and pyrroloquinoline quinone synthases.

### Plant growth and bacterial inoculation

Plant growth and bacterial inoculation involved the inoculation of diesel fuel-spiked soils with the isolated consortium in order to determine effects on plant growth and hydrocarbon change. The soil used for this experiment was “Primaster turf”. Primaster turf is made from a mixture of screened sand, soil, and composted organics. It is blended with a nitrogen-phosphorous-potassium fertiliser to promote root growth throughout the year. The soil texture was sand (88.6% sand, 6.1% silt and 5.3% clay), with 12.5% organic matter content measured by loss on ignition. The soil had a total nitrogen content of 0.15% and a pH of 7.1. The soil was sieved using a 2-mm sieve to remove large particles. Diesel fuel (C_10_-C_25_) from a Shell service station in Goettingen was added to the soil and homogenized following the methods of Eze, et al. ^17^ with some modifications. In brief, the soil was manually homogenized for an hour. This was followed by automatic homogenization using a portable soil mixing machine (Güde Model GRW 1400) for 30 minutes. Gas chromatography-mass spectrometry analysis revealed that the resulting total petroleum hydrocarbons concentration in the diesel fuel-contaminated soil was 4.59 g/kg. Viable seeds of *Medicago sativa* L. were placed in pots (3 seeds per pot) containing 150 g of diesel fuel contaminated soils.

The contaminated soil was treated with the following: (1) *M. sativa;* (2) Consortium; (3) *M. sativa* + consortium. An unplanted and uninoculated soil served as the control. Since the goal of the study was to assess the effectiveness of each treatment for hydrocarbon degradation, the soil used for the entire experiment was the diesel fuel-contaminated soil (4.59 g/kg) as described above. The microbial consortium used was harvested from the culture by centrifugation at 4000 × g for 10 min, washed twice in mineral medium and concentrated to OD_600_ = 1.800. For the *M. sativa* + consortium treatment, the cells (at OD_600_ = 1.800) were inoculated to the base of one-week-old *Medicago sativa* L. seedlings at the depth of 1.5 cm below-ground level. The whole experiment was performed in triplicate, and pots were watered with 90 mL sterile water every three days for the first two weeks. After that, the planted pots were watered with 90 mL sterile water every two days to compensate for the water needs of *M. sativa* plants. After 60 days, the plants were harvested, washed under tap water, oven-dried at 70 °C until constant weights were achieved, and then their dry biomass weights were obtained.

### Organic geochemical analysis of biodegradation

At the end of the experimental period, soils in each pot were homogenized as described earlier. Soils were freeze dried and 1 g of the ground freeze-dried soil was further homogenized with a small amount of sodium sulfate (Na_2_SO_4_) and transferred into a Teflon microwave digestion vessel for hydrocarbon analysis. The samples were solvent extracted twice with fresh 2.5 mL n-hexane each in a microwave device (Mars Xpress, CEM; 1600W, 100 °C, 20 min). For reference, 2.5 μL diesel fuel (density = 0.82 g/mL) was dissolved in 5 mL *n*-hexane instead of 1 g soil sample. The extracts for each sample were combined into a 7 mL vial and topped to 5 mL with *n*-hexane. A 1 mL aliquot (20%) of each extract was pipetted into a 2 mL autosampler vial, and 20 μL n-icosane D42 [200 mg/L] was added as an internal quantification standard.

Gas chromatography-mass spectrometry (GC-MS) analyses of the samples were performed using a Thermo Scientific Trace 1300 Series GC coupled to a Thermo Scientific Quantum XLS Ultra MS. The GC capillary column used was a Phenomenex Zebron ZB–5MS (30 m, 0.1 μm film thickness, inner diameter 0.25 mm). Compounds were transferred splitless to the GC column at an injector temperature of 300 °C. Helium was used as the carrier gas at a flow rate of 1.5 mL/min. The GC temperature program was as follows: 80 °C (hold 1 min), 80 °C to 310 °C at 5 °C/min (hold 20 min). Electron ionization mass spectra were recorded at 70eV electron energy in full scan mode (mass range m/z 50–600, scan time 0.42 s). Peak areas were integrated using Thermo Xcalibur software version 2.2 (Thermo Fisher Scientific Inc., USA).

### Statistical analysis

All statistical analysis were performed using R ^63^. One-way analysis of variance (ANOVA) was used to compare the mean dry biomass of “*M. sativa”* and “*M. sativa* + consortium” treatments. Similarly, comparisons of soil residual hydrocarbon concentrations between treatments were made using one-way ANOVA, followed by Tukey’s all-pairwise comparisons. In all cases, the normality of variances was tested by the Shapiro-Wilk’s method ^64^, while homogeneity of variances was tested using Levene’s test ^65^. Differences were considered significant at *p* < 0.05. The *p* values were adjusted using the Holm method, as this approach offers a simple, yet uniformly powerful method to control family-wise error rate in multiple comparisons ^66,67^.

## Data Availability

Raw sequencing data has been deposited in the sequence read archive of the National Center for Biotechnology Information under BioProject number **PRJNA612814**. The accession numbers used in this study are SRR11310428 and SRR11310429.

## Author contributions

Conceptualization and design: MOE, GCH, SCG and RD. Planning and implementation: MOE, SCG and RD. Experiments and bioinformatics analyses: MOE. Geochemical analysis: MOE and VT. Writing – original draft: MOE. Writing – review and editing: VT, GCH, SCG and RD. Supervision: GCH, SCG and RD.

## Competing interests

The authors hereby declare no competing interests.

## Acknowledgements

The authors would like to thank the Commonwealth Government of Australia and the German Academic Exchange Service (DAAD) for supporting this research project by providing M.O.E. with an international Research Training Program (iRTP) Scholarship and DAAD Scholarship (Allocation Numbers: 2017561 and 91731339 respectively). This publication was supported financially by the Open Access Grant Program of the German Research Foundation (DFG) and the Open Access Publication Fund of the University of Goettingen. We also thank Dr. Anja Poehlein and Melanie Heinemann for the assistance during sequencing.

## Notes

### Competing Interest Statement

The authors have declared no competing interest.

