## Supplementary Materials for "Metagenomic insight into the plant growth-promoting potential of a diesel-degrading bacterial consortium for enhanced rhizoremediation application"

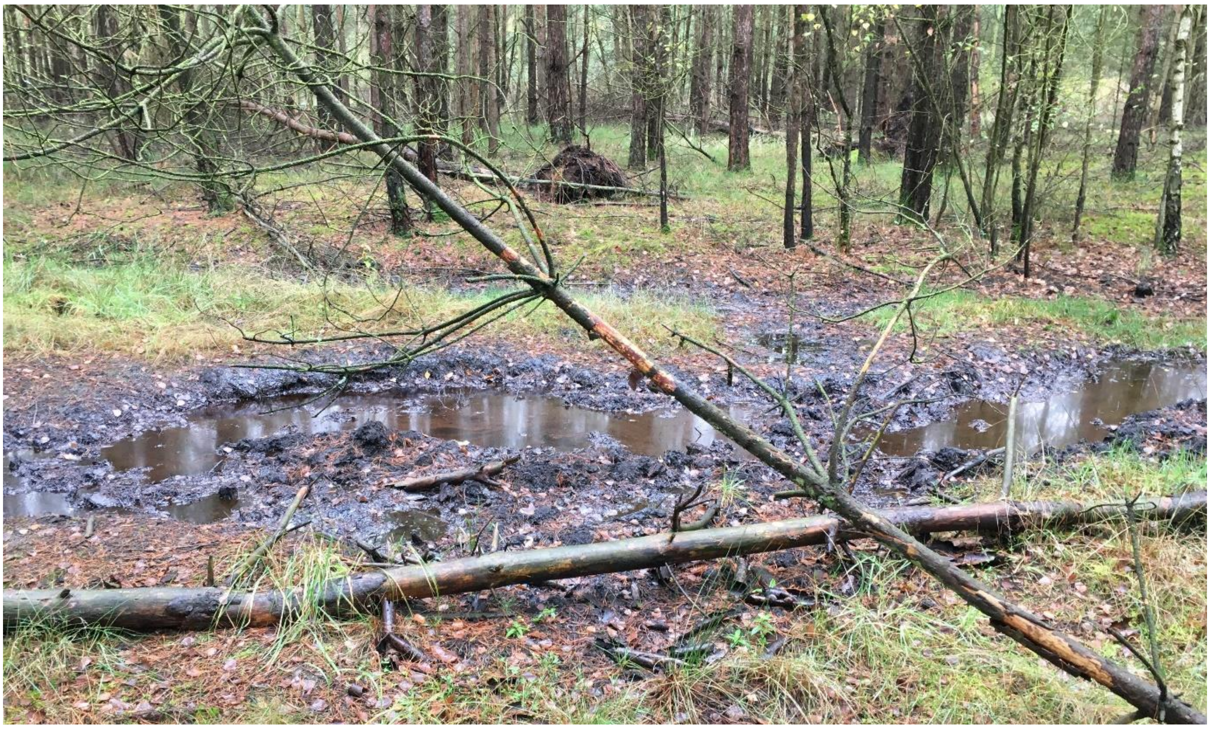

**Supplementary Figure S1**. Sampling site at Wietze, Germany (52°39’0’’N, 09°50’0’’E).

**Supplementary Table S1**. Genes involved in plant growth promotion, number of coding DNA sequences (CDSs), and their taxonomic classification.

| Group | Genes | KEGG Orthology | No. of CDSs | Taxa (Class) | Taxa (Genus) |
|---|---|---|---|---|---|
| | | | **Nitrogen fixation** | | |
| **Nitrogen uptake** | *nifA* | K02584 | 1 | Alphaproteobacteria | *Acidocella aminolytica* |
| | | K02584 | 4 | Acidobacteriia | *Acidobacterium capsulatum* |
| | | K02584 | 1 | Alphaproteobacteria | *Bradyrhizobium mercantei* |
| | | K02584 | 1 | Alphaproteobacteria | *Rhodobacter* |
| | | K02584 | 1 | Alphaproteobacteria | *Methylocella silvestris* |
| **Nitrogen uptake** | *por, nifJ* | K03737 | 2 | Acidobacteriia | *Acidobacterium capsulatum* |
| | | K03737 | 1 | Actinobacteria | *Mycobacterium malmoense* |
| | | K03737 | 2 | Alphaproteobacteria | *Asaia* sp. W19 |
| | | K03737 | 1 | Alphaproteobacteria | *Acetobacter senegalensis* |
| | | K03737 | 1 | Alphaproteobacteria | *Acetobacter cibinongensis* |
| **Nitrogen uptake** | *iscU, nifU* | K04488 | 1 | Acidobacteriia | *Acidobacterium capsulatum* |
| | | K04488 | 1 | Alphaproteobacteria | *Acidocella aminolytica* |
| **Nitrogen uptake** | *fixA, etfB* | K03521 | 1 | Alphaproteobacteria | *Elioraea* sp. YIM 72297 |
| | | K03521 | 1 | Alphaproteobacteria | unclassified *Rhizobiales* |
| | | K03521 | 1 | Acidobacteriia | *Acidobacterium capsulatum* |
| | | K03521 | 1 | Alphaproteobacteria | *Labrys okinawensis* |
| | | K03521 | 3 | Alphaproteobacteria | *Acidocella aminolytica* (2) and *Acidocella* (1) |
| | | K03521 | 1 | Alphaproteobacteria | *Acidiphilium multivorum* AIU301 |
| **Nitrogen uptake** | *fixB, etfA* | K03522 | 1 | Alphaproteobacteria | *Komagataeibacter* |
| | | K03522 | 1 | Alphaproteobacteria | unclassified *Rhizobiales* |
| | | K03522 | 1 | Acidobacteriia | *Acidobacterium capsulatum* |
| | | K03522 | 3 | Alphaproteobacteria | *Acidocella aminolytica* (2) and *Acidocella* (1) |
| | | K03522 | 1 | Alphaproteobacteria | *Acidiphilium multivorum* AIU301 |
| **Nitrogen uptake** | *fixL* | K14986 | 4 | Acidobacteriia | *Acidobacterium capsulatum* |
| | | K14986 | 2 | Alphaproteobacteria | *Rhodopseudomonas palustris* |
| | | K14986 | 4 | Alphaproteobacteria | *Acidocella aminolytica* (2) and *Acidocella* (2) |
| | | K14986 | 1 | Alphaproteobacteria | unclassified *Rhizobiales* |
| **Nitrogen uptake** | *fixJ* | K14987 | 1 | Alphaproteobacteria | unclassified Alphaproteobacteria |
| | | K14987 | 2 | Alphaproteobacteria | *Acidocella aminolytica* (1) and *Acidocella* (1) |
| | | K14987 | 2 | Alphaproteobacteria | unclassified *Rhizobiales* |
| **Nitrogen uptake** | *fixK* | K15861 | 4 | Alphaproteobacteria | *Acidocella aminolytica* (1) and *Acidocella* (3) |

| | | | | | |
|---|---|---|---|---|---|
| | | K15861 | 1 | Alphaproteobacteria | *Rhizomicrobium* sp. SCGC AG-212-E05 |
| | | K15861 | 1 | Alphaproteobacteria | *Roseomonas rosea* |
| | | K15861 | 1 | Alphaproteobacteria | *Acidiphilium* |
| | | **Subtotal:** | **55** | | |
| **Phosphate solubilization** | | | | | |
| **Phosphate solubilization** | *acpS* | K00997 | 4 | Alphaproteobacteria | *Acidocella aminolytica* (2) and *Acidocella* (2) |
| **Phosphate solubilization** | *acpP* | K02078 | 1 | Alphaproteobacteria | *Skermanella stibiiresistens* |
| | | K02078 | 2 | Acidobacteriia | *Acidobacterium capsulatum* |
| | | K02078 | 3 | Alphaproteobacteria | *Acidocella aminolytica* (2) and *Acidocella* (1) |
| | | K02078 | 1 | Betaproteobacteria | *Hydrogenophaga* |
| **Phosphate solubilization** | *phoA, phoB* | K01077 | 1 | Acidobacteriia | *Acidobacterium capsulatum* |
| **Phosphate solubilization** | *serB, PSPH* | K01079 | 2 | Acidobacteriia | *Acidobacterium capsulatum* |
| | | K01079 | 3 | Alphaproteobacteria | *Acidocella aminolytica* (2) and *Acidocella* (1) |
| **Phosphate solubilization** | *cysQ* | K01082 | 6 | Alphaproteobacteria | *Acidocella aminolytica* (3) and *Acidocella* (3) |
| | | K01082 | 1 | Alphaproteobacteria | *Acidiphilium* |
| | | K01082 | 1 | Alphaproteobacteria | *Sphingomonas* sp. YR710 |
| **Phosphate solubilization** | *otsB* | K01087 | 1 | Alphaproteobacteria | unclassified *Rhizobiales* |
| | | K01087 | 7 | Alphaproteobacteria | *Acidocella aminolytica* (4) and *Acidocella* (3) |
| | | K01087 | 2 | Acidobacteriia | *Acidobacterium capsulatum* |
| | | K01087 | 1 | Betaproteobacteria | *Paraburkholderia* sp. BL23I1N1 |
| **Phosphate solubilization** | *E3.1.3.16* | K01090 | 3 | Acidobacteriia | *Acidobacterium capsulatum* |
| | | K01090 | 6 | Alphaproteobacteria | *Acidocella aminolytica* (4) and *Acidocella* (2) |
| | | K01090 | 1 | Chitinophagia | *Niastella* |
| **Phosphate solubilization** | *gph* | K01091 | 1 | Acidobacteriia | *Acidobacterium capsulatum* |
| | | K01091 | 1 | Verrucomicrobiae | *Pedosphaera parvula* |
| | | K01091 | 5 | Alphaproteobacteria | *Acidocella aminolytica* (4) and *Acidocella* (1) |
| **Phosphate solubilization** | *suhB* | K01092 | 2 | Acidobacteriia | *Acidobacterium capsulatum* |
| | | K01092 | 8 | Alphaproteobacteria | *Acidocella aminolytica* (4) and *Acidocella* (4) |
| **Phosphate solubilization** | *appA* | K01093 | 1 | Acidobacteriia | *Acidobacterium capsulatum* |
| **Phosphate solubilization** | *pgpA* | K01095 | 1 | Acidobacteriia | *Acidobacterium capsulatum* |
| | | K01095 | 3 | Alphaproteobacteria | *Acidocella aminolytica* (2) and *Acidocella* (1) |

| | | | | | |
|---|---|---|---|---|---|
| **Phosphate solubilization** | *plc* | K01114 | 4 | Acidobacteriia | *Acidobacterium capsulatum* |
| | | K01114 | 5 | Alphaproteobacteria | *Acidocella aminolytica* |
| | | K01114 | 1 | Betaproteobacteria | *Paraburkholderia* sp. PDC91 |
| | | K01114 | 1 | Alphaproteobacteria | unclassified *Rhizobiales* |
| **Phosphate solubilization** | *cobC, phpB* | K02226 | 9 | Alphaproteobacteria | *Acidocella aminolytica* (6); *Acidocella facilis* (1) and *Acidocella* (2) |
| **Phosphate solubilization** | *cbiB, cobD* | K02227 | 1 | Alphaproteobacteria | *Roseovarius* sp. A-2 |
| | | K02227 | 1 | Alphaproteobacteria | *Acidocella aminolytica* |
| **Phosphate solubilization** | *glpR* | K02444 | 7 | Alphaproteobacteria | *Acidocella aminolytica* (5) and *Acidocella* (2) |
| | | K02444 | 1 | Alphaproteobacteria | *Methylovirgula* sp. 4M-Z18 |
| **Phosphate solubilization** | *glpX* | K02446 | 1 | Alphaproteobacteria | *Acidiphilium* |
| | | **Subtotal:** | **99** | | |
| **Zinc solubilization** | | | | | |
| **Zinc solubilization** | *pqqL* | K07263 | 3 | Acidobacteriia | *Acidobacterium capsulatum* |
| | | K07263 | 2 | Alphaproteobacteria | *Acidocella aminolytica* (1) and *Acidocella facilis* (1) |
| | | **Subtotal:** | **5** | | |
| **Siderophore transport** | | | | | |
| **Siderophore transport** | *entS* | K08225 | 1 | Acidobacteriia | *Acidobacterium capsulatum* |
| | | K08225 | 1 | Betaproteobacteria | unclassified *Burkholderiales* |
| | | K08225 | 1 | Alphaproteobacteria | *Acidocella* |
| | | **Subtotal:** | **3** | | |
| **Indoleacetic acid (IAA) synthesis** | | | | | |
| **IAA synthesis** | *iaaH* | K21801 | 1 | Alphaproteobacteria | *Hyphomicrobium* sp. 99 |
| | | K21801 | 1 | Alphaproteobacteria | *Mesorhizobium* |
| | | **Subtotal:** | **2** | | |
| **Pyrroloquinoline quinone synthesis** | | | | | |
| **PQQ synthesis** | *pqqB* | K06136 | 1 | Alphaproteobacteria | *Acidiphilium* |
| | | K06136 | 2 | Alphaproteobacteria | *Acidocella aminolytica* (1) and *Acidocella* (1) |
| **PQQ synthesis** | *pqqC* | K06137 | 1 | Acidobacteriia | *Acidobacterium capsulatum* |
| | | K06137 | 2 | Alphaproteobacteria | *Acidocella aminolytica* (1) *and Acidocella* (1) |
| **PQQ synthesis** | *pqqD* | K06138 | 3 | Alphaproteobacteria | *Acidocella aminolytica* (2) and *Acidocella* (1) |
| **PQQ synthesis** | *pqqE* | K06139 | 1 | Alphaproteobacteria | *Bosea* sp. 117 |
| | | K06139 | 1 | Acidobacteriia | *Acidobacterium capsulatum* |
| | | K06139 | 2 | Alphaproteobacteria | *Acidocella aminolytica* (1) and *Acidocella facilis* (1) |
| | | **Subtotal:** | **13** | | |
| | | **TOTAL:** | **177** | | |
